# Role of *cis*, *trans*, and inbreeding effects on meiotic recombination in *Saccharomyces cerevisiae*

**DOI:** 10.1101/361428

**Authors:** Xavier Raffoux, Mickael Bourge, Fabrice Dumas, Olivier C. Martin, Matthieu Falque

## Abstract

Meiotic recombination is a major driver of genome evolution by creating new genetic combinations. To probe the factors driving variability of meiotic recombination, we used a high-throughput method to measure recombination rates in 26 *S. cerevisiae* strains from different geographic origins and habitats. Fourteen intervals were monitored for each strain, covering chromosomes VI and XI entirely, and part of chromosome I. We found an average number of crossovers per chromosome ranging between 1.0 and 9.5 across strains (“domesticated” or not), which is higher than the average between 0.5 and 1.5 found in most organisms. In the different intervals analyzed, recombination showed up to 9-fold variation across strains but global recombination landscapes along chromosomes varied less. We also built an incomplete diallel experiment to measure recombination rates in one region of chromosome XI in 10 different crosses involving five parental strains. Our overall results indicate that recombination rate is increasingly positively correlated with sequence similarity between homologs (i) in DSB rich regions within intervals, (ii) in entire intervals, and (iii) at the whole genome scale. Therefore, these correlations cannot be explained by *cis*-effects only. In addition, by using a quantitative genetics analysis, we identified an inbreeding effect that reduces recombination rate in homozygous genotypes while other interaction effects (specific combining ability) or additive effects (general combining ability) are found to be weak. Finally, we measured significant crossover interference in some strains, and interference intensity was positively correlated with crossover number.

**Author Summary:** Meiosis is a key process for sexually reproducing organisms by producing gametes with a halved set of genetic material. An essential step of meiosis is the formation of crossovers which are reciprocal exchanges of genetic material between chromosomes inherited from both parents. Crossovers ensure proper chromosome segregation and thus viable gametes. They also create novel genetic diversity which contributes to evolution and permits genetic improvement of agriculturally important species. Most living organisms produce between one and three crossovers per chromosome, and tight regulatory mechanisms control the number of crossovers and their distribution along chromosomes. In spite of their potential importance for biotechnological applications, such mechanisms are still poorly understood.

Using a high throughput method based on fluorescent markers, we investigated the diversity of recombination in the budding yeast Saccharomyces cerevisiae. We observed up to 9-fold differences in numbers of crossovers across hybrids obtained by crossing different strains with a common tester, and this variation was correlated with the degree of DNA sequence similarity between homologous chromosomes. By also investigating homozygotes, we conclude that on the one hand too much sequence divergence impairs recombination in distantly-related hybrids, and on the other hand complete homozygosity is also associated with lower numbers of crossovers.

## Introduction

In sexually reproducing organisms, meiosis is a particular type of cell division producing gametes that contain half of the somatic genetic material. Meiotic recombination is a major driver of genome dynamics and evolution in sexually reproducing organisms because it generates new allelic combinations that can be subject to natural selection. The number of crossing-over events and their positions along the chromosomes are tightly regulated, but the mechanisms involved are still not well understood. Getting more insights into the regulation of recombination rate and crossover distribution would be beneficial for many fields of fundamental and applied genetics, in particular to improve the efficiency of plant breeding [1]. Meiotic recombination starts by programmed DNA double-strand breaks throughout the genome. DSB repair occurs using the homologous chromosome as template, which in turn allows recognition and pairing of the homologous chromosomes. DSB repair is achieved by different pathways, leading to either crossovers (COs), that are reciprocal exchanges of genetic material, or non-crossovers (NCOs), for which genetic change is limited to a small DNA segment around the break. In most organisms, the distribution of DSBs and COs is not homogeneous along chromosomes. At a fine scale (a few kilobases), they are clustered in regions called hotspots, as has been shown for instance in *S. cerevisiae* [2–4] and in humans (60% of COs lying in such hotspots; [5]). In *S. cerevisiae*, 84% of CO hotspots overlap with gene promoters [6]. At the chromosome scale, large “hot” regions showing high CO rates alternate with colder regions. The peri-centromeric regions of the chromosomes are “cold” in most organisms (S. *cerevisiae* [6], *Arabidopsis thaliana* [7], maize [8] and tomato [9]). DSBs occur usually in open chromatin regions. In human and mice, many DSB hotspots occur in DNA sequences targeted by the histone H3K4 methyltransferase PRDM9 [10]. In *S. cerevisiae*, the SET1 complex deposits histone H3K4 methylation at the positions of future DSB regions, where the SPP1 protein [11] makes a link between H3K4me3 and SPO11 which in turn generates DSBs [12]. In maize and *A. thaliana*, CO-rich regions are correlated with low DNA methylation [13,14] and low transposable element content [15,16]. In *S. cerevisiae* and *A. thaliana*, DSB hot spots co-localize with transcriptionally active regions, especially promoters [2,17]. In *S. cerevisiae*, approximately 40% of DSBs are repaired to form COs, the other DSBs being repaired as NCOs or using the sister chromatid as template [18,19]. The ratio between CO and DSB numbers can be regulated at two levels during DSB repair: (1) by driving repair to the homologous chromosomes *vs* the sister chromatids, and (2) by choosing the repair pathway leading to the formation of COs *vs* NCOs. [20–23]. CO numbers vary at both inter- and intra-species levels. However, for 76% of the species studied (fungi, animals, and plants), the number of COs per bivalent ranges from 1 to 3 (see review [24]). This low variation in CO numbers across species suggests selective constraints keeping recombination levels within a certain range. The presence of at least one CO per homologous pair can be explained by the need to ensure correct chromosome segregation during the first meiotic division. Concerning the upper limit, possible selective pressures might prevent too many DSBs from becoming COs [24]. But this hypothesis remains speculative, especially because some species such as *S. cerevisiae* and *S. pombe* can produce 10 or more COs per bivalent. Furthermore it was recently shown that in *A. thaliana* the number of COs can be increased about nine-fold without perturbing chromosome segregation [25]. CO numbers also vary at the intra specific level [8,26–28], though this variation is generally smaller than between species. In *S. cerevisiae*, using four parental strains [26], it was observed that (1) CO hotspots as well as cold spots, are highly conserved among crosses, (2) the number of COs per meiosis varies from 48 to 64.5, and (3) the recombination rate varies up to 60% between strains in some intervals. Relatively few studies have investigated the variation of meiotic recombination rate at a broad level within one species. In the present work, we characterized the intra-specific diversity of recombination rate in a large part of the *S. cerevisiae* genome. To do so, we used a high-throughput method to measure crossover rates [29] in diploids obtained by crossing a SK1 strain to 26 strains taken from a core-collection of *S. cerevisiae* strains (Supp Tab 1). To measure recombination, each strain of the core collection was crossed with eight SK1 testers carrying three different fluorescent markers (mCherry, yECerulean, and Venus, respectively denoted RFP, CFP, and YFP) at different chromosomal locations (see Materials and Methods). In the resulting diploids, we measured the recombination rate and CO interference based on 14 genomic segments covering chromosomes VI and XI, and part of chromosome I. Our results show up to 2.5 fold differences in recombination rate when considering all pooled intervals and up to 9-fold differences in some intervals. Our dataset indicates also a clear positive correlation between CO numbers and genome wide sequence similarity between homologs in the hybrids, and thus a negative correlation between recombination and observed heterozygosity. However, concomitantly, the correlation was weaker when using sequence similarity within the interval where recombination is measured. To obtain further insights, five strains were intercrossed in an incomplete diallel design (among the fifteen possible parental combinations, only ten crosses produced diploids able to sporulate). The recombination rate of these ten diploids was then analyzed in one interval of chromosome XI. Altogether, we find (1) that sequence similarity between homologs (and thus heterozygosity) plays a major role in the observed variation of recombination rate, and (2) that homozygosity lowers recombination, a phenomenon that can be thought of as an inbreeding depression.

## Results

### Sporulation, spore viability and recombination rate

Because of the large genetic diversity explored in this work, we first assessed the correct progress of meiosis using sporulation rate and spore viability as proxies. When crossing all strain of the collection (see Materials and Methods; [67]) with SK1, sporulation rates at the plateau (always reached after 10 days on the sporulation medium; Supp Fig 1) ranged from 14% to 85% across hybrids with a continuous variation, the maximum being reached for the SK1×SK1 diploid which is completely homozygous (Supp Fig 2A). Spore viability ranged from 1.5 to 85 %, the hybrids from strains UWOPS03_461_4, UWOPS05_217_3, UWOPS05_227_2 (Malaysian wild strains), and YS9 (Asian baking strain) producing almost no viable spores (Supp Fig 2B). Such low viability may denote abnormalities in the meiotic process, e.g. associated with possible chromosomal rearrangements. Therefore we discarded these four strains. Spore viability was not correlated with the sporulation rate (*p*-value=0.16) indicating that these two biological processes are relatively independent. Finally, the average recombination rate over the eight testers was significantly positively correlated with spore viability (*r*^2^=0.49 *p*-*value*=7.2×l0^−5^).

### Wide diversity of recombination rate in the collection

When pooling the information obtained from all intervals of the eight testers, we obtained global recombination rates ranging from 0.20 cM/kbp to 0.51 cM/kbp across the 22 hybrids tested. The highest value corresponds to the SK1 × SK1 hybrid (Fig 1). Recombination rates averaged over hybrids varied significantly between chromosome I (0.47 cM/kbp), chromosome VI (0.39 cM/kbp), and chromosome XI (0.30 cM/kbp) (Tukey’s HSD test: *p*-value < 10^−7^; Supp Fig 3), and between individual testers as well as between individual intervals delimited by fluorescent markers (Supp Fig 3; Supp Tab 2). The patterns of recombination rate along chromosomes were significantly different between hybrids for some intervals, but all hybrids showed the same decreasing recombination rate tendency in the vicinity of centromere regions except for chromosome I for which there is a strong DSB hotspot in the interval containing the centromere (Fig 2). For each interval, the ratio between the most and least recombining hybrids ranged from 1.8 to 9.5. Note that the SK1 × K1 diploid had the highest recombination rate only for intervals two and ten. Analyses of variance revealed significant effects of hybrids, intervals, and hybrid × interval interactions on recombination rate (*p*-value < 2.2×l0^−16^ for each effect). Further, we observed a significant effect of the geographic origin on the global (eight testers pooled) recombination rate of the hybrid (ANOVA *p*-value = 0.009). Specifically, pairwise significant differences were observed between African and American origins (Tukey’s HSD test: *p*-value = 0.049). Genome-wide sequence-based phylogenetic groups [31] also showed a significant association with recombination rate (ANOVA *p*-value = 3.9×10^−6^). Specifically, pairwise significant differences were observed between the West-African group and all other groups (Tukey’s HSD test: *p*-values < 10^−4^). Because adaptation to a changing environment can drive evolution towards higher recombination rate [32], we analyzed hybrids of strains grown solely in laboratory habitat (supposed to be a stable environment). Surprisingly, they had significantly higher recombination rates than the strains coming from all other types of habitat (Tukey’s HSD test: *p*-values < 10^−4^) (Supp Fig 4).

**Figure 1:**
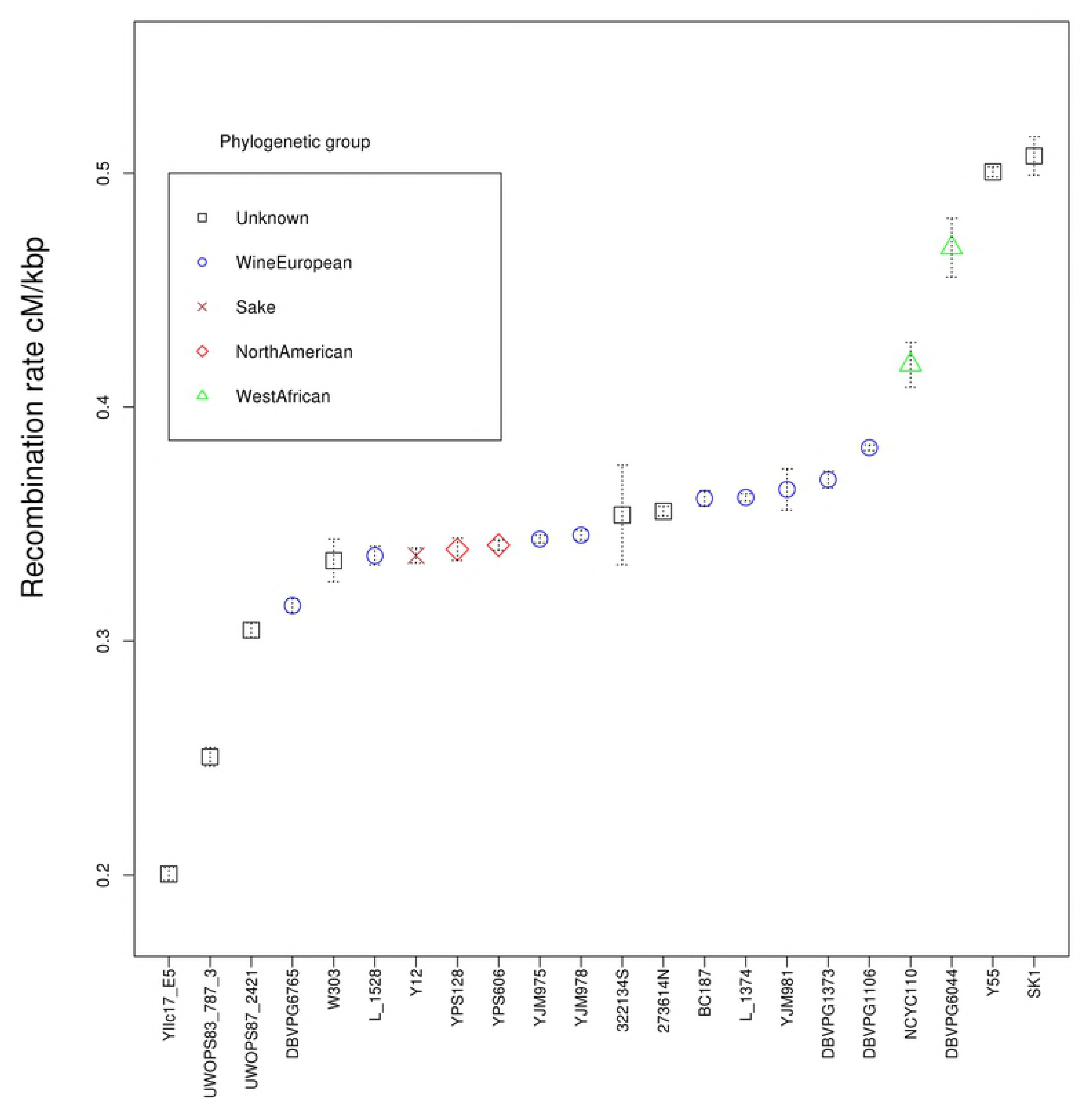
Average recombination rate over the 14 intervals for each strain of the collection crossed to SK1 testers. Symbols and colors refer to the phylogenetic group of the strains. Error bars indicate 95% confidence intervals based on four biological replicates.

**Figure 2:**
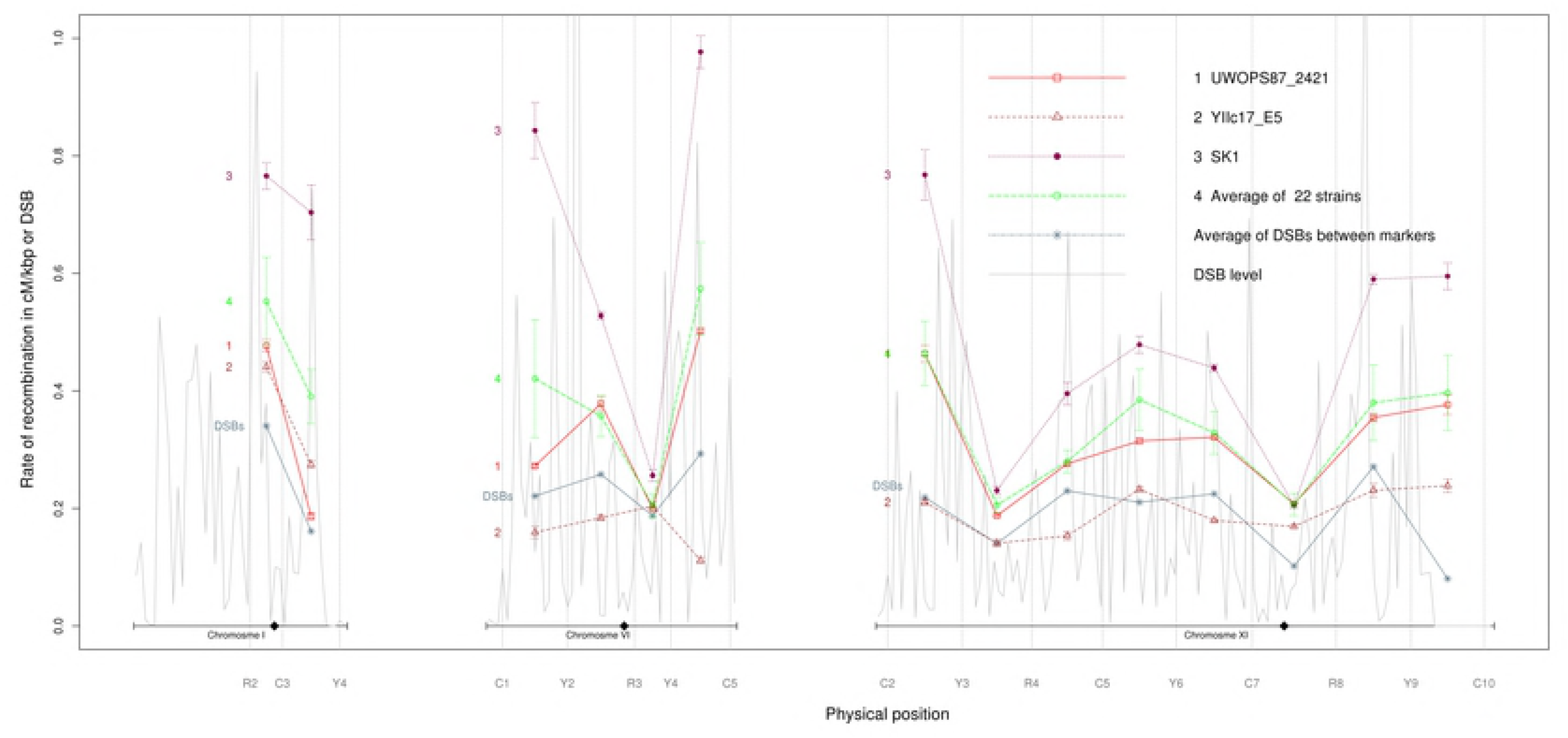
Recombination rates in cM/kbp along chromosomes for hybrids between SK1 and strains UWOPS87_2421, YIIc17_E5, and SK1. Error bars indicate 95% confidence intervals based on four biological replicates. Also shown: (1) frequency of double strand breaks per base (Pan *et al*. 2011) between markers along chromosomes I, VI, and XI, and (2) DSB level: average number of DSBs per 5kb window. Vertical dashed lines indicate the positions of fluorescent markers. Horizontal lines at the bottom indicate chromosome boundaries and diamonds show centromere positions.

### Relationship between recombination rate and DSB levels

For all hybrids, recombination rates and DSB patterns showed a positive correlation except in the region between markers Y2 and R3 of chromosome VI, and in the region between Y9 and C10 of chromosome XI (Fig 2). Patterns of recombination rate along chromosomes and average SK1 DSB levels (data from Pan *et al*. [4]) were both low near the centromere, except for chromosome I (Fig 2).

### High levels of heterozygosity reduces recombination

To investigate the correlation of recombination rate with sequence similarity between homologous chromosomes (which is one minus the observed heterozygosity) across the different hybrids, we considered successively five scales of sequence similarity: the pool of all intervals studied, the pool of all intervals on each chromosome, each interval separately, DSB-rich regions within each interval, and 30Kb regions surrounding each interval (see Materials and Methods). We found a significant positive correlation between average recombination rate and sequence similarity when pooling all intervals (r^2^=0.43 *p*-*value*=9×l0^−4^) (Fig 3), as well as when pooling intervals for each chromosome (r^2^>0.2 *p*-*value*<0.04) (Supp Fig 5). The three chromosomes investigated thus seem to have similar correlations. When considering the 14 intervals separately, we found significant positive correlations between sequence similarity and recombination rate for nine of them. Analysis of sequences flanking these 14 intervals on both sides showed that only five intervals gave significant positive correlations (Supp Tab 3). Finally, focusing on sequence similarity within DSB-rich regions in these 14 intervals, nine intervals showed significant positive correlations (Supp Tab 3). Interestingly, the correlation between recombination rate and sequence similarity in DSBs rich regions is weaker than when considering the whole sequence spanned by intervals, showing that CO number is not mainly controlled by local sequence similarity at the sites of DSBs repair. Similarly, the correlation between the recombination rate and sequence similarity within the interval studied is weaker than when considering genome wide sequence similarity, which points to the existence of significant *trans* effects that may be more important than *cis* effects for controlling CO number.

**Figure 3:**
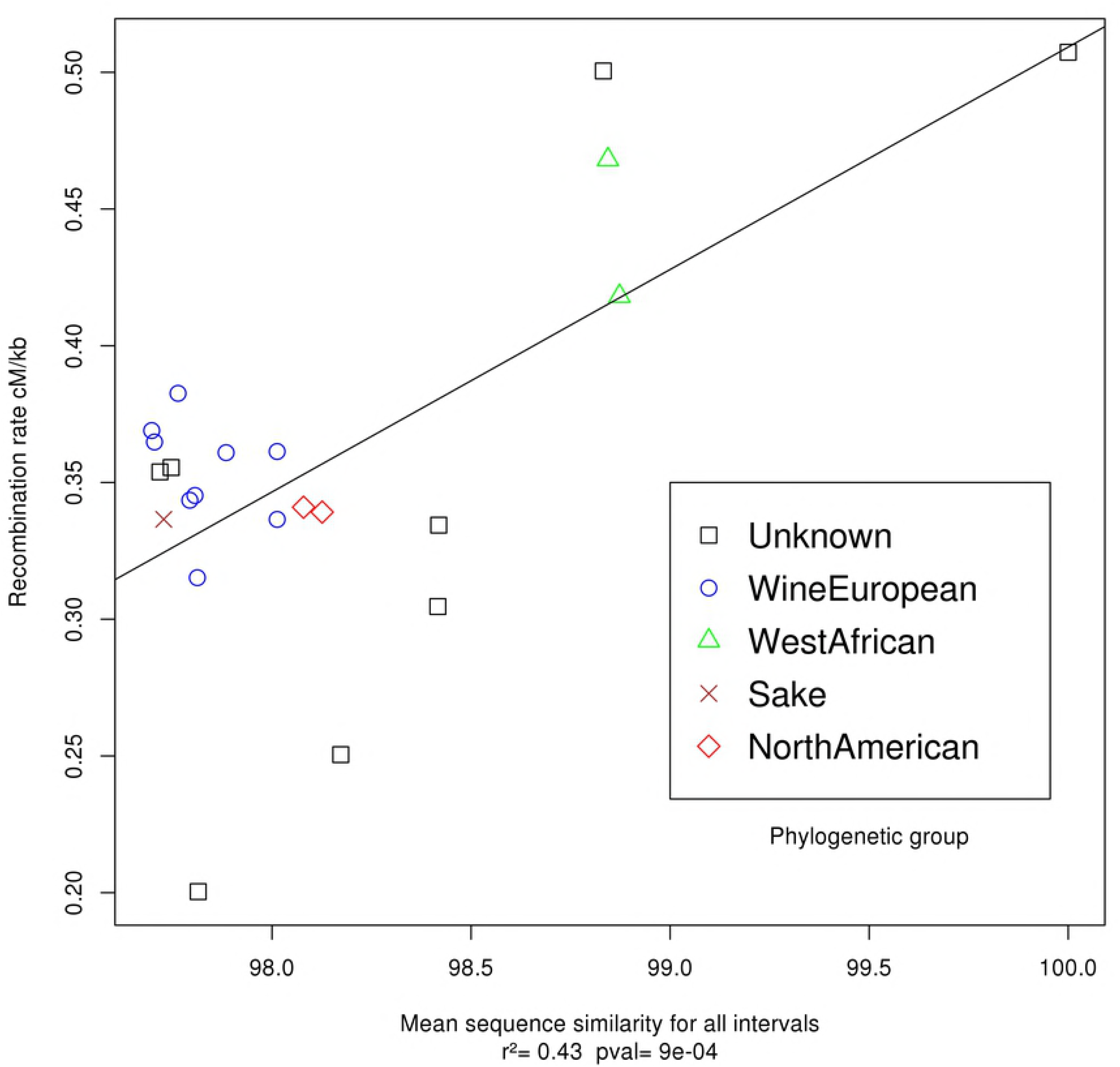
Correlation between sequence similarity when pooling all intervals and the mean recombination rate of hybrids. x-axis: score of sequence similarity (see Materials & Methods), y-axis: for each strain of the collection, average of the eight recombination rates of the hybrids obtained by crossing the strain with the eight testers. The legend indicates the geographic origin of the strains.

### Crossover interference analysis

To quantitatively compare interference strength across strains and chromosomal regions, we used the *ν* parameter of the gamma model [33], inferred for each pair of adjacent intervals (corresponding to one tester) from its coefficient of coincidence (CoC) and its two recombination fractions measured for a strain × tester combination (see Materials & Methods and Supp Methods). As in our previous study [29], we discarded the tester SK1-XI-R1C2Y3 from interference analyses because its first interval is too small (5,557 bp). When pooling the information given by the seven testers, we obtained *ν* values ranging from 0.54 to 1.53 across hybrids (Supp Fig 6). Most hybrids show either no interference (*ν* ≈1) or positive interference (*ν* >1). However, the two strains YIIc17_E5 and UWOPS83_787_3, which also have the lowest genome-wide recombination rates, display negative interference (*ν* <1). Interference patterns along chromosomes were also significantly different between some hybrids (Supp Fig 7). We found significant effects of hybrid, tester, and interaction hybrid × tester on interference strength (ANOVA *p*-value < 2.2×l0^−16^). Interference strength and average recombination rate were positively correlated across the seven testers (Supp Fig 8), even when discarding the two outlier strains YIIc17_E5 and UWOPS83_787_3 from the data (r2=0.56, *p*-*val*=10^−4^).

## Inbreeding reduces recombination

To obtain further insights on the control of recombination rate, we measured the genetic length of interval Y9C10 on chromosome XI for 10 hybrids obtained by crossing five parental strains in an incomplete diallel experiment (See Fig 4). As above, the recombination rates in this diallel experiment showed a significant correlation with sequence similarity between homologs (*p*-*value*= 7×10^−10^, *r*^2^=0.4). Recombination rate is a quantitative trait displaying genetic diversity (Fig 1). As such, it may be controlled by several types of mechanisms involving QTLs possibly interacting with each other. These QTLs may have two kinds of effects: (1) additive effects of individual alleles, which sum up in the hybrid, referred to as general combining ability (GCA; [34]), and (2) interaction effects between alleles either at the same locus (including dominance, over-dominance, and inbreeding) or between alleles at different loci (epistasis), referred to as specific combining ability (SCA; [34]). The effect of heterozygosity on recombination rate may be considered as a particular type of SCA because there is no additive effect associated with individual sequences and the recombination rate depends on each pair of homologous sequences. Therefore we used the pairwise sequence similarity as a quantitative explicative variable in our diallel analysis, thereby distinguishing sequence similarity effects from other interaction effects. We then used the Hierarchical Generalized Linear Model below, considering sequence similarity between homologs as a *fixed* effect, and GCA, SCA, and inbreeding as *random* effects. Specifically, the statistical model sets

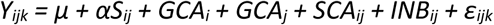

*Y_ijk_*: Genetic distance (in cM), measured in the hybrid formed by crossing strain *i* and strain *j* for the replicate *k*

*μ*: Intercept (in cM)

*α*: Coefficient associated with sequence similarity effect (in cM per percent of similarity)

*S_ij:_* Percentage of sequence similarity between strain *i* and strain *j* (See Materials and Methods)

*GCA_i_*: General combining ability (in cM) of strain *i*

*GCA_j:_* General combining ability (in cM) of strain *j*

*SCA_ij:_* Specific combining ability (in cM) of the hybrid obtained by crossing strain *i* and strain *j* when *i*≠*j, set to 0 when i=j*, calculated as *Y_ij_* - 1/2 (*Y*_*i*._ + *Y*_.*j*_) - *μ*

*INB_ij:_* Inbreeding effect when *i* = *j* (in cM), calculated as *Y_ii_* - *Y*_*i*._ - *μ*

*ε_ijk:_*Residual variance

**Figure 4:**
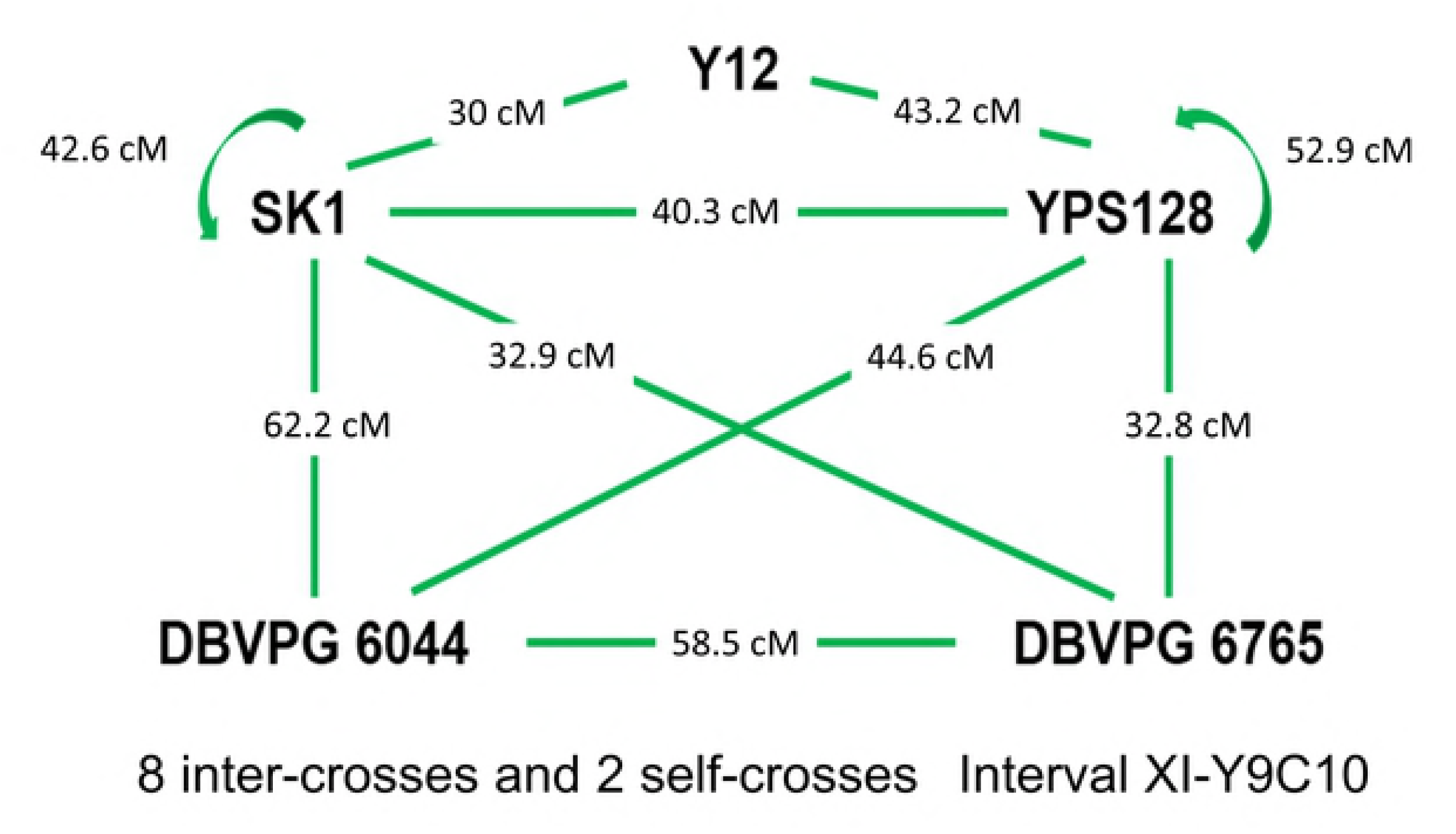
Hybrids obtained by crossing five parental strains. Each arrow represents a cross and corresponding numbers indicate the genetic distance in centiMorgan measured in the interval.

To estimate the parameters, we used the R hglm package [35,36] as described by [37] and [38], which uses a Bayesian approach to fit hierarchical generalized linear models. We found that the Akaike information criterion (H. Akaike, 1973) decreased when adding factors one after the other, indicating that all parameters of the model are relevant. We further checked that there was a strong significant correlation between experimental and predicted phenotypic values (*r*^2^=0.78 *p*-value=4.5×l0^−45^) (Supp Fig 9).

The results of the diallel analysis are given in Supp Fig 10. Values of effects are relative to the intercept *μ* which would be the phenotypic value obtained if all effects were null. The GCA results showed that strain DBVPG6044 had a significantly higher GCA value (+8.5cM from the intercept value) than the four other strains (between -3.4 and 0cM), which were not very different from each other. SCA results showed some differences between parental combinations (ranging from -3cM to +4.5cM) but the effects remained limited. In the cases of SK1 and YPS128, for which we could measure the recombination rates in the homozygous diploids, we observed strong inbreeding effects, in effect depressing the recombination rates in a major way (*INB*=-36,7cM for SK1 and *INB*=-23.3cM for YPS128). Finally, the estimated effects of sequence similarity (*αS*) ranged from 0 to +44.4cM across hybrids, the two highest values corresponding to homozygous diploids. Thus in our experiment, hybrids from distantly related strains show that heterozygosity decreases recombination rate, but we also see from the two homozygotes having negative inbreeding effects, that high levels of homozygosity might also decrease recombination rate. Altogether, *αS* and *INB* effects were much stronger than other effects, suggesting that sequence similarity may be the strongest factor driving the genetic diversity of recombination rate within *S. cerevisiae* strains.

## Discussion

### Intraspecific diversity of recombination

When the level of divergence between homologous chromosomes is too high, DSBs cannot be repaired through the homologous recombination pathway but may be repaired through the mismatch repair pathway, leading to aneuploidy and loss of spore viability [39,40]. In our study, we thus discarded the hybrids showing strong spore viability defects, to keep only those which are relevant for studying homologous recombination. We observed recombination rates (averaged across the eight testers) between 0.20 and 0.51 cM/kbp, which is consistent with previous results in budding yeast genome-wide analyses: 0.4cM/kbp [26] (see their fig 2), 0.61 cM/kbp [6], or from 0.29 to 0.63 cM/kbp for chromosome VII left arm [41]. Across our 22 strains, we obtained an average 2.55-fold variation of recombination rate, that can be compared to the Cubillos *et al*. [26] observation of a 4-fold variation between crosses of four genetically distant *S. cerevisiae* strains. In maize, close to 30% variation was measured in genome wide CO numbers between 23 [8] and 25 [28] hybrids, based on genetic mapping. The present study also indicates that the recombination landscape along chromosomes is different across strains: the ratio between the most and least recombining hybrids ranged from 1.8 to 9.5 depending on the interval. In maize [28] an average 2.9-fold variation in CO number was reported between 25 hybrids, some intervals showing up to 30-fold differences. Such high levels of variation of recombination rate across intervals suggest that determinants affect recombination in close-by locations (*cis* effects). Finally, we observed significantly different recombination rates depending on the habitat of the parental strains, but there is no indication that strains living in changing environments may have evolved higher recombination rates to adapt more easily, as previously hypothesized [32,42–44]. In fact, creating more genetic combinations can play positive roles for adaptation but can also have deleterious effects by breaking up established favorable arrangements.

### Intraspecific diversity of crossover interference

Through our measurements of coefficient of coincidence (CoC) and interferene strength (v), we observed positive CO interference for most testers and hybrids, which is in accordance with previous studies reporting interference in *S. cerevisiae* [41,45–47].

Two hybrids however, YIIc17_E5 × SK1 and UWOPS83_787_3 × SK1, showed negative interference. These two hybrids are among those with the lowest recombination rate, sporulation rate, and spore viability of the collection. In such crosses between distantly related parents, negative interference can be justified *a posteriori* as being due to meiotic defects.

Specifically, if two homologs simply do not pair in some fraction of meioses, CO events will be statistically positively correlated. In a similar vein, if homologs stochastically pair only along part of their length, COs will be restricted to those paired regions and thus in effect they will be subject to clustering. Both situations result in apparent negative interference even if there is positive interference between crossovers for each meiosis. This is in direct analogy with what was observed in the *Arabidopsis axr1* mutant [48].

We also observed significant variation of interference across hybrids. To our knowledge, intraspecific diversity of interference strength had never been assessed before in *S. cerevisiae*, but in maize, Bauer *et al*. [8] reported significant differences among 23 hybrids based on the gamma model. We also measured variations of interference intensity along and between chromosomes, as already reported in *S. cerevisiae* [6,49] and in *Arabidopsis* [50]. Our results showed significant positive correlations (averaged across seven testers) between recombination rate and interference strength, whereas in maize, Bauer *et al*. [8] reported a significant negative correlation. It is commonly hypothesized that interference reduces CO number while ensuring the obligatory CO [51]. Indeed, there seems to be selective pressure against too many COs, although the reasons are unclear [24]. The maize results of Bauer *et al.* [8] are in accordance with this hypothesis, whereas our results in yeast are not. An explanation may come from the fact that in maize, each meiocyte undergoes almost 500 DSBs which produce about 20 COs [52,53], whereas in *S. cerevisiae*, 40% of DSBs leads to the formation of COs [54].The DSB/CO ratio is then about 25 in maize to be contrasted with 2.5 in *S. cerevisiae*. So in the context of selective pressure against too many COs, CO regulation through interference will be much more efficient in maize than in yeast, which might explain the difference between our results in yeast and results in maize [8].

### Genetic control of recombination rate

#### Effect of heterozygosity and homozygosity on recombination rate

Considering all intervals pooled, the recombination rates in our study showed a significant positive correlation with sequence similarity between the two parents of the hybrid. This result is in accordance with previous studies showing that heterozygosity can have an inhibitory effect on homologous recombination in yeast [55] or in *A. thaliana* [56]. In our study, sequence similarity in DSB rich regions did not explain recombination rate better than sequence similarity in whole intervals, suggesting that the sequence similarity in the region of strand invasion is probably not the main determinant of DSB commitment into CO *vs* NCO. However, we used DSBs pattern obtained from a homozygous SK1 strain, whereas our hybrids are heterozygous between SK1 and other strains, so DSBs landscape may be different in our hybrids although they have one haplotype in common. Elsewhere our analysis of sequence similarity in regions flanking the 14 intervals on both sides showed a significant positive correlation with the recombination rate within the interval in five cases (four of which also being significant when considering sequence similarity within the intervals). This suggests that those flanking regions might carry some of the determinants of the positive correlation between sequence similarity and recombination rate. Similarly, results on *A. thaliana* [57] showed that the presence of a heterozygous interval next to a homozygous region leads to more COs in the heterozygous region and less in the homozygous one. At a larger scale, we observed that the genome wide correlation between recombination rate and sequence similarity is stronger than when focusing on individual chromosomes, and even more than when focusing on individual intervals. This points to the presence of *trans* acting factors modulating CO formation, in addition to possible *cis* effects. Our results altogether suggest that heterozygosity alone is not sufficient to explain the variation observed in CO numbers and positions across hybrids, as previously reported [8,58–60,60]. CO control may also depend on other factors such as structural differences between homologous genomes that can (1) inhibit CO formation as observed in *A. thaliana* [61], or (2) modify CO frequency as suggested in maize [8,58]. Beyond sequence-related effects, recombination can also be modulated by epigenetics factors, as observed in centromeric regions [14,16,62], and by environmental conditions [63–65].

#### Dissecting parental effects on recombination

Our diallel experiment also showed a significant positive effect of sequence similarity on recombination rate. Together with the correlation observed in our diversity experiment, this confirms that heterozygosity is a major determinant of the intraspecific genetic diversity of recombination rate in *S. cerevisiae* hybrids. Since this determinant is defined pairwise rather than in terms of individual sequences, its effect has no additive component and may be considered as overdominance. This is illustrated by comparing recombination rates of heterozygous *vs* homozygous crosses involving the parental strains SK1 and YPS128 (Fig 4; sup Fig 11): crossover numbers measured in SK1 × YPS128 were significantly lower than in both SK1 × SK1 and YPS128 × YPS128 crosses, reflecting overdominance due to sequence divergence. But in fact, the quantitative analysis of the diallel experiment revealed that this apparent sequence similarity effect comes from the combined effects of *αS* and *INB* which represent respectively the negative effect of strong heterozygosity on recombination and *also* the negative effect of inbreeding in perfect homozygotes on recombination (*αS=+44 cM, INB*=-36.7cM for SK1 × SK1 and *INB*=-23.3cM for YPS128 × YPS128) (Supp Tab 4). Accordingly, the highest values of recombination in the diallel experiment do not correspond to homozygous diploids but to the heterozygous hybrid SK1 × DBVPG6044. This may be explained by the fact that SK1 and DBVPG6044 may be genetically close enough to allow high recombination rates (*αS*=23,35cM) but different enough to escape inbreeding effects. It would be interesting to extend our experiment to more closely related strains to investigate more precisely such inbreeding effect. It is usually assumed that inbreeding depression is due to recessive deleterious mutations [66], and this is expected to be particularly true in outcrossing species which did not purge such mutations. In the case of *S. cerevisiae*, the HO gene can lead to mating type switch [67] which may favor inbreeding, but the level of outcrossing in natural *S. cerevisiae* populations remains unknown [68,69].

## Materials and Methods

### Biological material

The collection of 26 *S. cerevisiae* strains used in this study comes from the Saccharomyces Genome Resequencing Project (SGRP; [67]), and strains were kindly provided by F. Cubillos, Universidad de Santiago de Chile, Santiago, Chile. These strains were collected from various geographical areas and types of habitats (Supp Tab 1). The eight SK1 tri-fluorescent testers strains hereafter referred to as “testers” (SK1-I-R2C3Y4, SK1-VI-C1Y2R3, SK1-VI-R3Y4C5, SK1-XI-R1C2Y3, SK1-XI-Y3R4C5, SK1-XI-R4C5Y6, SK1-XI-Y6C7R8, and SK1-XI-R8Y9C10) used to measure recombination are described in [29]. Each of them contains three reporter genes distant by around 30 centiMorgans on a same chromosome, coding for three different fluorescent proteins that can be detected in flow cytometry. That nice feature allowed us to use tri-fluorescent testers rather than bi-fluorescent ones, speeding up the process of measuring recombination rates; as a bonus, we also obtained measures of genetic interference since we were able to detect the presence of double recombinants.

### Sporulation efficiency

Each of the 26 Mat a strains of the collection was crossed with the Mat α tester SK1-XI-R1C2Y3 to produce a hybrid diploid and spores as described in [29]. At days 1-2-3-4-7-8-9-10-11 of incubation on solid SPOR medium (2.5% yeast extract, 1% glucose, 10% potassium acetate) at 30°C, cells were picked up and resuspended in 10μL H_2_O on a microscope slide. Tetrads and vegetative cells were counted at 1000X magnification.

### Spore viability

At day 10 of the sporulation efficiency experiment, we scraped one quadrant of each of the 26 Petri dishes and prepared spores as described in [29] for FACS sorting. We selected events corresponding to the size of spores using a gate in the side scatter (SSC)-Height-Log *vs* forward scatter (FSC)-Height-Log graph (Summit software, Beckman Coulter, USA), then we discarded events containing more than one cell using a gate in the SSC-Height-Log *vs* SSC-Area-Log graph (see Materials and Methods in [29]). One spore per well was distributed in two 96-wells plates containing 100μL solid YPD medium. After 48 hours incubation at 30°C, we counted the number of wells in which a colony had grown. In rare cases, two colonies were observed in the same well and these events were discarded from further analyses. Thus, 192 spores were analyzed per condition.

### Recombination rate and interference measurements on the collection

As it was technically impossible handle all strain and all testers in the same experiment, we worked with each tester, one at a time. Thus, for one given experiment, each of the 26 Mat a strains of the collection was deposited with one Mat α tri fluorescent tester strain on solid YPD medium and incubated one night at 30°C to produce diploid cells and then transferred to sporulation medium (SPOR). To capture possible variation due to environmental heterogeneity, the experiment was designed in the following manner: (1) for each cross, four Petri dishes were placed at different positions in the incubator to provide four replicates, and (2) in each experiments, the control (Y12 Mat a) × (SK1-VI-Y3R4C5 Mat α) diploid was added. After ten days at 30°C, tetrads were picked up by scraping one quarter of the Petri dish surface, and spores were then isolated as described in [29]. The spore suspensions were analyzed with a MoFlo ASTRIOS flow cytometer (Beckman Coulter, USA) and the associated software Summit. Vegetative cells were filtered out based on SSC and FSC as described above, and then the fluorescence intensity was analyzed for each spore in the mCherry, yECerulean, and Venus channels (excitation at 561, 405, and 488 nm respectively, emission at 614/20, 448/59, and 526/52 nm respectively). To quantify recombination rates and coefficients of coincidence (CoC), we used the mathematical model given in [29] to take into account the fact that fluorescence can be extinguished at a low rate (see Supp Methods; Supp Fig 12). As recombination rate values of the (Y12 Mat a) × (SK1-VI-Y3R4C5 Mat α) control sample didn’t show significant variation between the eight experiments corresponding to the eight testers (ANOVA p-value = 0.99), results were normalized using this control as a standard (see Supp Methods). The coefficient of coincidence (CoC) for a pair of intervals is defined as the ratio between the *experimental* frequency of double recombinants and its *theoretical* frequency in the absence of interference. Absence of interference means that recombination events in the two intervals are independent, and thus the theoretical frequency is simply the product of each interval’s recombination rate. Since CoC values strongly depend on recombination rate in the two intervals, we cannot compare CoC values across different strains or testers. We thus used a simulation approach to map the correspondence between CoC and the parameter *ν* of the gamma model [32] for each strain / tester combination. First, it was necessary to simulate the relationship between recombination fraction and number of crossovers for each value of *ν* (see examples in Supp Fig 13), and then the relationship between CoC and *ν* (see examples in Supp Fig 14; see details in Supp Methods). The parameter *ν* is a quantitative measurement of interference strength, it’s value is 1 in the absence of interference, greater than 1 in the presence of positive interference, and lower than 1 in the presence of negative interference. In the gamma model framework, this parameter does not depend on recombination rate and thus its values may be compared across strains and testers.

### Score of sequence similarity at different scales

Reference sequences of all strains studied come from the Saccharomyces Genome Resequencing Project (SGRP; [30,68]). The sequence similarity percentage between homologous genomes was calculated at different scales: (1) genome-wide, (2) in the whole chromosome carrying the considered markers, (3) in the interval surrounded by the two markers analyzed, (4) within that interval, but focusing only on the DSBs-rich regions defined as 300bp regions for which Pan et al. found at least 100 Spo11-associated oligo reads [4], and (5) in the 30kb regions surrounding the interval. Similarity percentages were calculated both-ways, using the SK1 sequence as query blasted against the other parent as subject, and the reciprocal analysis using the SK1 sequence as subject and the other parent as query. Motivated by what occurs during the repair of meiotic double strand breaks, for each pair of sequences considered, the query sequence was sliced in 200bp windows sliding with a 50bp step. Only windows which did not contain any “N” in their sequence (92.6 % of the cases, sd = 7.9 %) were considered. For each window, we calculated the sequence similarity percentage as the fraction of identical nucleotides in the first High-Scoring-segment Pair multiplied by its length and divided by the size of the window (200) and multiplied by 100. We then took the average percentage of similarity for all windows within the region considered, calculated both ways. These computations were carried out using R scripts calling standalone BLAST+ [70]. Blast was preferred to sequence alignment software because it is much quicker and complete alignments were not necessary here.

## Acknowledgements

The authors thank Christine Dillmann and Marianyela Petrizzelli for their help in analyzing the diallele experiment, as well as Valérie Borde, Monique Bolotin-Fukuhara and Denise Zickler for helpful comments on the manuscript. The present work has benefited from the Cytometry/Electronic Microscopy/Light Microscopy facility of Imagerie-Gif (http://www.i2bc.paris-saclay.fr), a member of IBiSA (http://www.ibisa.net).

## Supporting Information Legends

**Raffoux_et_al_SUPP_FIG1.pdf:** this file contains Supplementary Figure 1

**Raffoux_et_al_SUPP_FIGURES.pdf:** This file contains all supplementary figures except Supplementary Figure 1

**Raffoux_et_al_SUPP_TABLES.pdf:** this file contains all Supplementary Tables

**Raffoux_et_al_SUPP_METHODS.pdf:** this file contains additional explanations on the methods used

